# Optimised production of disulfide-bonded fungal effectors in *E. coli* using CyDisCo and FunCyDisCo co-expression approaches

**DOI:** 10.1101/2021.08.31.458447

**Authors:** Daniel S. Yu, Megan A. Outram, Emma Crean, Ashley Smith, Yi-Chang Sung, Reynaldi Darma, Xizhe Sun, Lisong Ma, David A. Jones, Peter S. Solomon, Simon J. Williams

## Abstract

Effectors are a key part of the arsenal of plant pathogenic fungi and promote pathogen virulence and disease. Effectors typically lack sequence similarity to proteins with known functional domains and motifs, limiting our ability to predict their functions and understand how they are recognised by plant hosts. As a result, cross-disciplinary approaches involving structural biology and protein biochemistry are often required to decipher and better characterise effector function. These approaches are reliant on high yields of relatively pure protein, which often requires protein production using a heterologous expression system. For some effectors, establishing an efficient production system can be difficult, particularly those that require multiple disulfide bonds to achieve their naturally folded structure. Here, we describe the use of a co-expression system within the heterologous host *E. coli* termed CyDisCo (cytoplasmic disulfide bond formation in *E. coli*) to produce disulfide bonded fungal effectors. We demonstrate that CyDisCo and a naturalised co-expression approach termed FunCyDisCo (Fungi-CyDisCo) can significantly improve the production yields of numerous disulfide bonded effectors from diverse fungal pathogens. The ability to produce large quantities of functional recombinant protein has facilitated functional studies and crystallisation of several of these reported fungal effectors. We suggest this approach could be useful when investigating the function and recognition of a broad range of disulfide-bond containing effectors.

## Introduction

Fungal pathogen infections are a leading cause of yield losses in many economically important crops. During infection and colonisation of their plant host, fungal pathogens utilise small, secreted virulence proteins, known as effectors, to promote disease (Stergiopoulos and de Wit 2009). Characterised effectors have been implicated in functions that include the targeting and disruption of plant defences and nutrient acquisition from the host (Selin et al. 2016). Effectors can also be recognised by plant receptors, which activate defence pathways leading to plant immunity (Dodds and Rathjen 2010).

Fungal pathogens utilise 10-1000s of effectors during colonisation of plant hosts (Oliveira-Garcia and Valent 2015). Understanding how these effectors function is often challenging. Many effectors have low sequence similarity to proteins with known functional domains or motifs, preventing reliable functional predictions based on sequence alone. The most informative effector function studies often require multi-discipline approaches.

We are interested in understanding the structure and function of effectors from multiple plant-pathogenic fungi. Many of these are Kex2-processed pro-domain (K2PP) effectors, which include cysteine-rich effectors with thiol groups of the cysteine sidechain involved in disulfide bond formation (Outram et al. 2021). To study disulfide-bond containing effectors, we have sought to develop tools to enhance protein production in *Escherichia coli* (Zhang et al. 2017; Outram et al. 2021). To this end, we (and others), have had success using the specialised strain of *E. coli*, SHuffle® (New England Biolabs, Ipswich, Massachusetts, United States) (Maqbool et al. 2015; Zhang et al. 2017; De la Concepcion et al. 2018; Outram et al. 2021). SHuffle is engineered to address unfavourable redox potential in the cytoplasm through disruption of the glutaredoxin and thioredoxin pathways, and expression of a cytoplasmic version of the disulfide bond isomerase protein, DsbC, which normally localises to the periplasm (Fig. 1A and B). These manipulations have been shown to improve production of correctly-folded disulfide-bonded proteins (Lobstein et al. 2012). We have subsequently utilised small solubility tags to further enhance disulfide-rich effector yields in <u>SHuffle</u> (Outram et al. 2021). Despite these advancements, the yields obtained for many of our effectors of interest have remained low and inadequate for structural and biochemical studies.

**Fig. 1.**
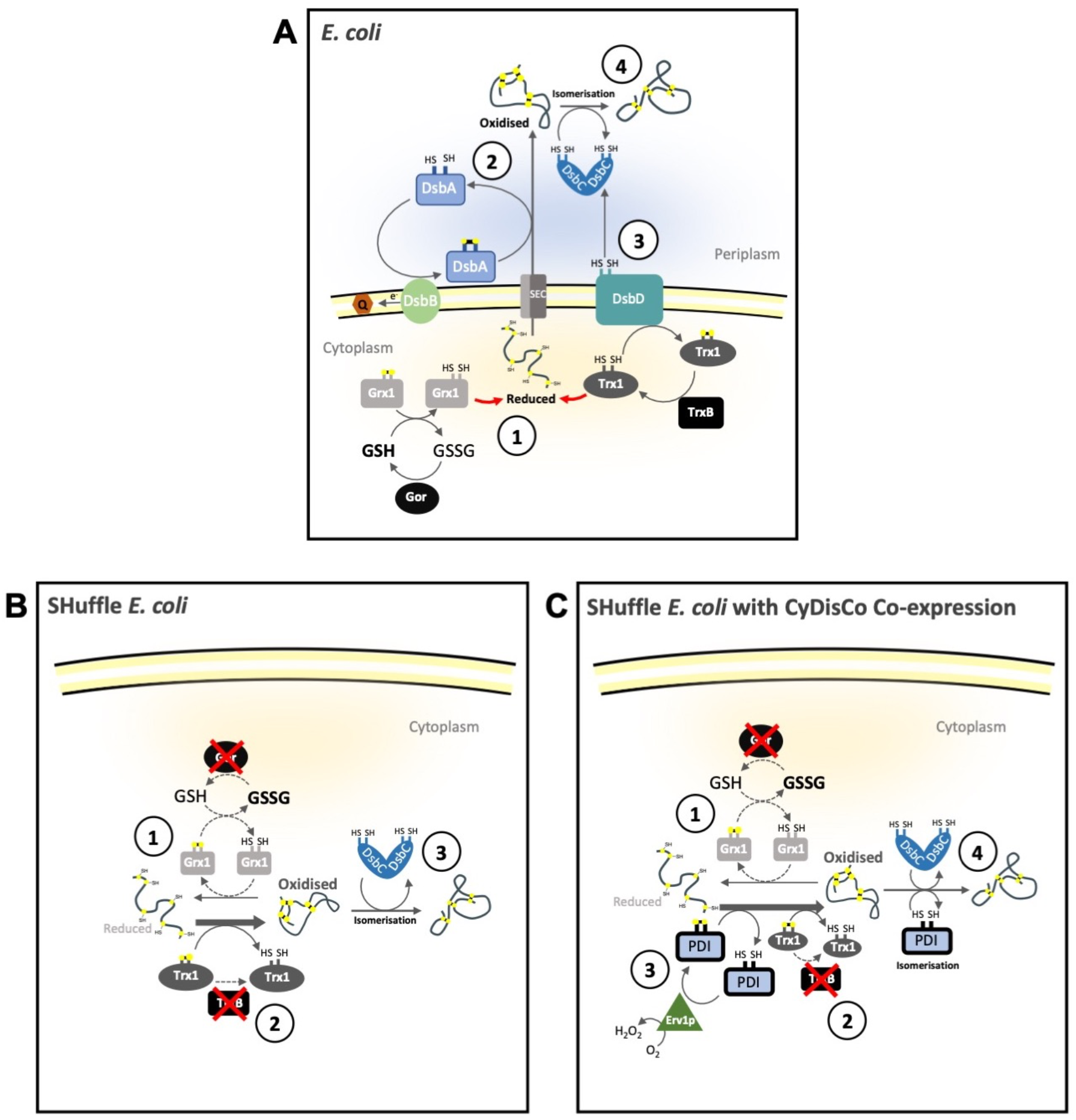
Disulfide bond formation in *Escherichia coli* expression systems. **(A)** In wild-type *E. coli*, proteins are produced in the cytoplasm in a reduced state. 1: The cytoplasm is a reducing environment. Contributing factors include the high reduced glutathione (GSH): oxidised glutathione (GSSG) ratio maintained by glutathione reductase (Gor). Excess GSH reduces glutaredoxin-1 (Grx1), which can then reduce nascent proteins. Proteins in the cytoplasm can also be reduced by thioredoxin-1 (Trx1), which maintains its reducing power by thioredoxin reductase (TrxB). 2: Newly translated proteins are transported out of the cytoplasm into the periplasm where they are oxidised by DsbA. The oxidative power of DsbA is regenerated by DsbB. Electrons are accepted by ubiquinone (Q) and carried to the electron transport chain. 3: Oxidised proteins may be incorrectly disulfide-bonded but can be isomerised by DsbC. For isomerisation to occur, DsbC needs to be in a reduced or hemi-reduced state which is maintained by DsbD. The redox state of DsbD is reset by cytoplasmic Trx1. 4: Once DsbC is in a reduced state, it can isomerise the disulfide bonds on the oxidised protein resulting in correct disulfide-bonding. **(B)** In SHuffle *E. coli* (Lobstein et al. 2012), the redox state of the cytoplasm is altered to be more oxidising. 1: The cytoplasm of SHuffle has a lower GSH: GSSG ratio due to the Gor knockout weakening the reduction pathway. 2: TrxB knockout prevents the reduction of Trx1, which is usually maintained to reset the redox state of DsbD. In conjunction with a weaker reduction pathway, the higher proportion of oxidised Trx1 strengthens the oxidation pathway and newly translated proteins can be oxidised in the cytoplasm. 3: Newly oxidised proteins in the cytoplasm may be incorrectly disulfide-bonded. SHuffle *E. coli* is engineered to cytoplasmically express DsbC, which can isomerise oxidised proteins in the cytoplasm. **(C)** CyDisCo co-expression in SHuffle *E. coli* further strengthens the oxidation pathway in the cytoplasm. 1: The Gor knockout and 2: TrxB knockout in Shuffle *E. coli* weakens the reduction pathway and strengthens the oxidation pathway, respectively. 3: CyDisCo co-expression of protein disulfide isomerase (PDI) readily oxidised newly translate proteins in the cytoplasm. The redox state of PDI is reset by the sulfhydryl oxidase, Erv1p, which generates *de novo* disulfide bonds by donating electrons on to O_2_. Erv1p can also oxidise proteins further strengthening the oxidation pathway. 4: Incorrectly disulfide-bonded proteins are isomerised by cytoplasmically expressed DsbC, from SHuffle, and PDI from CyDisCo.

To address this limitation, we have sought to further improve our production system. The emergence of synthetic biology and the molecular tools that support this discipline have seen an increased interest in co-expression of eukaryotic machinery and chaperones in *E. coli* to improve recombinant protein production (Zhou et al. 2018). This approach has also been developed to enhance production of disulfide-bonded proteins. In 2014, Matos and colleagues introduced the CyDisCo system (for **cy**toplasmic **dis**ulfide bond formation in *E*. ***co****li*) (Matos et al. 2014). CyDisCo involves co-expression, in *E. coli*, of a disulfide-bonded protein of interest with yeast mitochondrial sulfhydryl oxidase, Erv1p, and human protein disulfide isomerase (PDI) (Fig. 1C). To date, enhanced production of numerous disulfide-rich human proteins has been reported, including antibodies, human growth factor and perlecan (Matos et al. 2014; Gaciarz et al. 2016; Sohail et al. 2020). More recently, the CyDisCo system has been used to produce functional recombinant SARS-CoV-2 spike receptor binding domain (Prahlad et al. 2021).

Here, we demonstrate the utility of the CyDisCo co-expression system in combination with SHuffle *E. coli* to produce disulfide-rich fungal effectors. Using this system, seven out of eight effector candidates studied were successfully purified with higher yields (ranging from 2x to 29x) compared to SHuffle alone. We sought to naturalise the system further towards the production of fungal effectors by utilising a native PDI and sulfhydryl oxidase from *Fusarium oxysporum* f. sp. *lycopersici* (*Fol*). The naturalised system, termed FunCyDisCo, outperformed protein production using SHuffle alone and had varied, protein-dependent results, compared with CyDisCo. In our hands, the adoption of CyDisCo/FunCyDisCo has enabled the functional and structural investigation of numerous disulfide-rich effectors that could not otherwise be achieved. We suggest this approach could be broadly useful in the investigation of the function and recognition of a broad range of disulfide-bond containing effectors.

## Results

### CyDisCo facilitates the improved production of SIX6 proteins from *Fusarium oxysporum*

We have previously demonstrated that numerous *Fol* Secreted in Xylem (SIX) effectors can be produced using the *E. coli* strain SHuffle in combination with an N-terminal GB1 (protein GB1 domain) solubility tag (Outram et al. 2021). Nevertheless, yields remained relatively low for some effectors of interest, including SIX6 (∼0.3 mg/L of culture) and made structural studies difficult and laborious (Outram et al. 2021). To address this limitation, we employed CyDisCo, which involves co-expression of a sulfhydryl oxidase and PDI with the effector of interest in SHuffle *E. coli* (Fig. 1C). To understand the effectiveness of this approach, we performed side-by-side expression and purification of N-terminal 6xHis-GB1 tagged SIX6 (lacking the signal peptide) in SHuffle alone or in SHuffle with CyDisCo (Fig. 2A). GB1-SIX6 produced in SHuffle alone was highly expressed, however, most of the protein was insoluble. The total amount of SIX6, when co-expressed with CyDisCo was lower compared to SHuffle alone, however the total protein and soluble fraction (clarified lysate) were indistinguishable, suggesting improved solubility. GB1-SIX6 expressed with and without CyDisCo was subsequently purified from the soluble fractions using nickel affinity chromatography (Fig. 2B). The protein yields obtained for GB1-SIX6 with CyDisCo were greater than GB1-SIX6 produced in SHuffle alone, and of higher purity as determined by SDS-PAGE analysis (Fig. 2B). Purified protein obtained when SIX6 was expressed alone contained higher quantities of high molecular weight proteins consistent with the presence of heat-shock proteins compared to SIX6 co-expressed with CyDisCo. Heat-shock proteins typically assist in protein folding but can maintain associations with unfolded protein (Lesley et al. 2002). Their presence can indicate the existence of soluble aggregates of a protein of interest (in this case GB1-SIX6) and are typically a negative indicator of protein quality. We removed the GB1 fusion partner from SIX6 using 3C protease and used an additional nickel purification step to remove the GB1-tag, uncleaved protein and the 3C protease, prior to further purification by size exclusion chromatography (SEC). More mono-dispersed SIX6 protein was obtained when expressed with CyDisCo compared to SHuffle alone (Fig. 2C). The final yields were 1.5 mg/L for SIX6 co-expressed with CyDisCo, compared with 0.4 mg/L when expressed alone in this experiment. We consistently obtained higher final yields (ranging from 0.9 to 2.2 mg/L) for SIX6 co-expressed with CyDisCo compared to SIX6 expressed in SHuffle alone (from 0.2 to 0.5 mg/L) in three independent replicates of this experiment, highlighting the robustness and reproducibility of the co-expression approach. The quality of the purified SIX6 was analysed using intact protein mass spectrometry (MS) and circular dichroism (CD) spectropolarimetry, which revealed that the protein is disulfide bonded (four disulfides formed) and contains secondary structural elements dominated by β‐sheets (Supplementary Fig. S1 and S2).

**Fig. 2.**
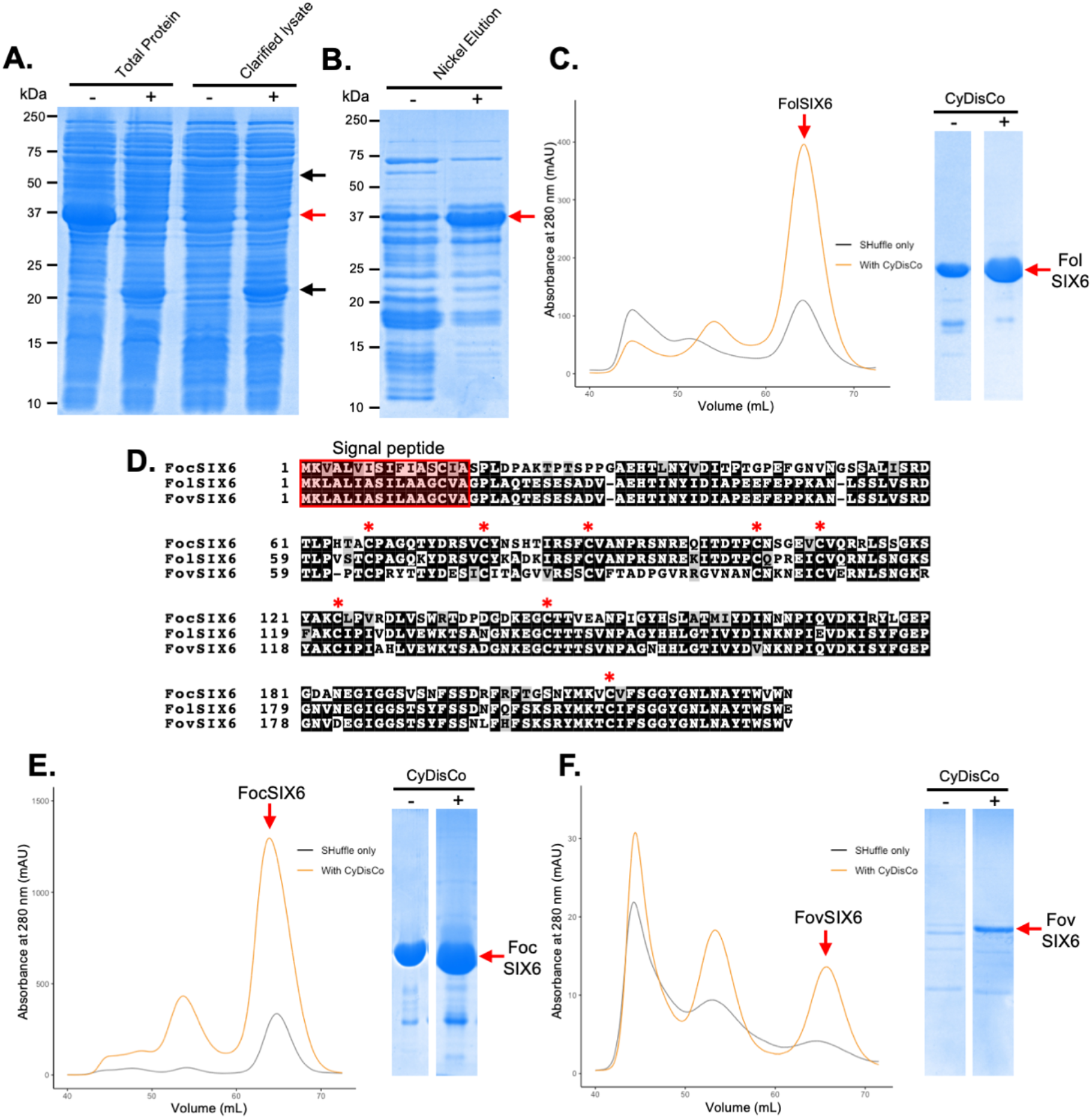
Recombinant SIX6 protein can be produced in SHuffle *E. coli* alone but yields are higher when co-expressed with CyDisCo. **(A)** Coomassie-stained SDS-PAGE gel showing total protein and soluble proteins from SHuffle *E. coli* expressing 6xHisGB1-FolSIX6 with (+) or without (-) CyDisCo co-expression. The red arrow points to the FolSIX6 (with N-terminal 6xHisGB1) protein band of ∼37 kDa. The black arrows point to the expression of soluble PDI (∼55 kDa) and Erv1p (∼21 kDa). **(B)** Coomassie-stained SDS-PAGE gel showing the proteins captured by immobilised metal affinity chromatography (IMAC) from SHuffle *E. coli* expressing 6xHisGB1-FolSIX6 with (+) or without (-) CyDisCo co-expression, with the red arrow indicating 6xHisGB1-FolSIX6. **(C)** Left panel: Size-exclusion chromatograms (SEC) of FolSIX6 protein produced by SHuffle *E. coli*, following cleavage by 3C protease to remove the N-terminal 6xHisGB1 fusion. Shown in orange is FolSIX6 produced in SHuffle with CyDisCo co-expression and in black is FolSIX6 produced in SHuffle alone. The red arrow points to the peak corresponding to FolSIX6. Right panel: Coomassie-stained SDS-PAGE gel showing equal volume loading of FolSIX6 protein (indicated by red arrow) expressed with (+) or without (-) CyDisCo corresponding to the protein peak from SEC. **(D)** Sequence alignment of FolSIX6 and two SIX6 homologues from *F. oxysporum* f. sp. *cubense* (Foc) and *F. oxysporum* f. sp. *vasinfectum* (Fov). The signal peptide is highlighted in red, as determined by SignalP (Almagro Armenteros et al. 2019). Conserved cysteine residues are marked with a red asterisk. Size-exclusion chromatogram and SDS-PAGE analysis for **(E)** FocSIX6 and **(F)** FovSIX6 produced with (orange trace) or without (black trace) CyDisCo co-expression, as presented in **(C)**.

Several homologues of SIX6 exist in different *forma specialis* of *F. oxysporum* ranging from 60–92% similarity to SIX6 from *Fol*. Notably, all homologues of SIX6 contain conserved cysteine residues (Fig. 2D). To validate the effectiveness of the CyDisCo system for improving protein yields of SIX6, homologues of SIX6 from *Foc* (*F. oxysporum* f. sp. *cubense*) and *Fov* (*F. oxysporum* f. sp. *vasinfectum*) were expressed with and without CyDisCo, and subsequently purified (Fig. 2E and F). The co-expression of CyDisCo improved the soluble protein yield by ∼5-fold for FocSIX6 from 0.6 mg/L to 3.1 mg/L, and ∼5-fold for FovSIX6 for FovSIX6 from 0.04 mg/L to 0.2 mg/L, consistent with the results for FolSIX6. Collectively, these data suggest that CyDisCo promotes improved disulfide-bond formation to boost yields of soluble correctly-folded SIX6 in *E. coli*.

### CyDisCo facilitates the improved production of an expanded set of disulfide-rich fungal effectors

Based on the success observed for SIX6 we wanted to test the utility of the CyDisCo system to produce different disulfide-rich fungal effectors. The effectors chosen include SIX1 (Avr3) and SIX4 (Avr1) from *Fol*, SnTox1 and SnTox3 from *Parastagonospora nodorum*, and NIP2.1 from *Rhynchosporium commune* (Supplementary Table S1). Most of these effectors could only be produced in low yields from SHuffle *E. coli* despite the addition of fusion partners (Outram et al. 2021). SIX1, SIX4 and SnTox3 were expressed with an N-terminal 6xHisGB1 tag, but GB1 was not included for SnTox1 and NIP2.1 as the tag was a similar size to the proteins of interest leading to complications during downstream analysis. Proteins expressed in Shuffle *E. coli* alone or with CyDisCo were expressed and purified (side-by-side) using the same approach described for SIX6 (details in methods) and the final mono-dispersed SEC elution fractions were compared (Fig. 3). In most cases, we observed an increase in final yields associated with co-expression with CyDisCo with a ∼29-fold improvement for SIX1 resulting in a yield of 4.3 mg/L, ∼6-fold improvement for SIX4 with a final yield of 2.4 mg/L, ∼3-fold improvement for SnTox1 with a final yield of 1.5 mg/L and ∼2.5-fold improvement for NIP2.1 with a final yield of 0.15 mg/L. SnTox3 was the only protein that did not show any obvious improvement in yield when co-expressed with CyDisCo. Collectively, this demonstrates a general effectiveness of the CyDisCo co-expression system in improving the yield of multiple disulfide-rich effectors across different fungal species, with the degree of improvement being protein specific.

**Fig. 3.**
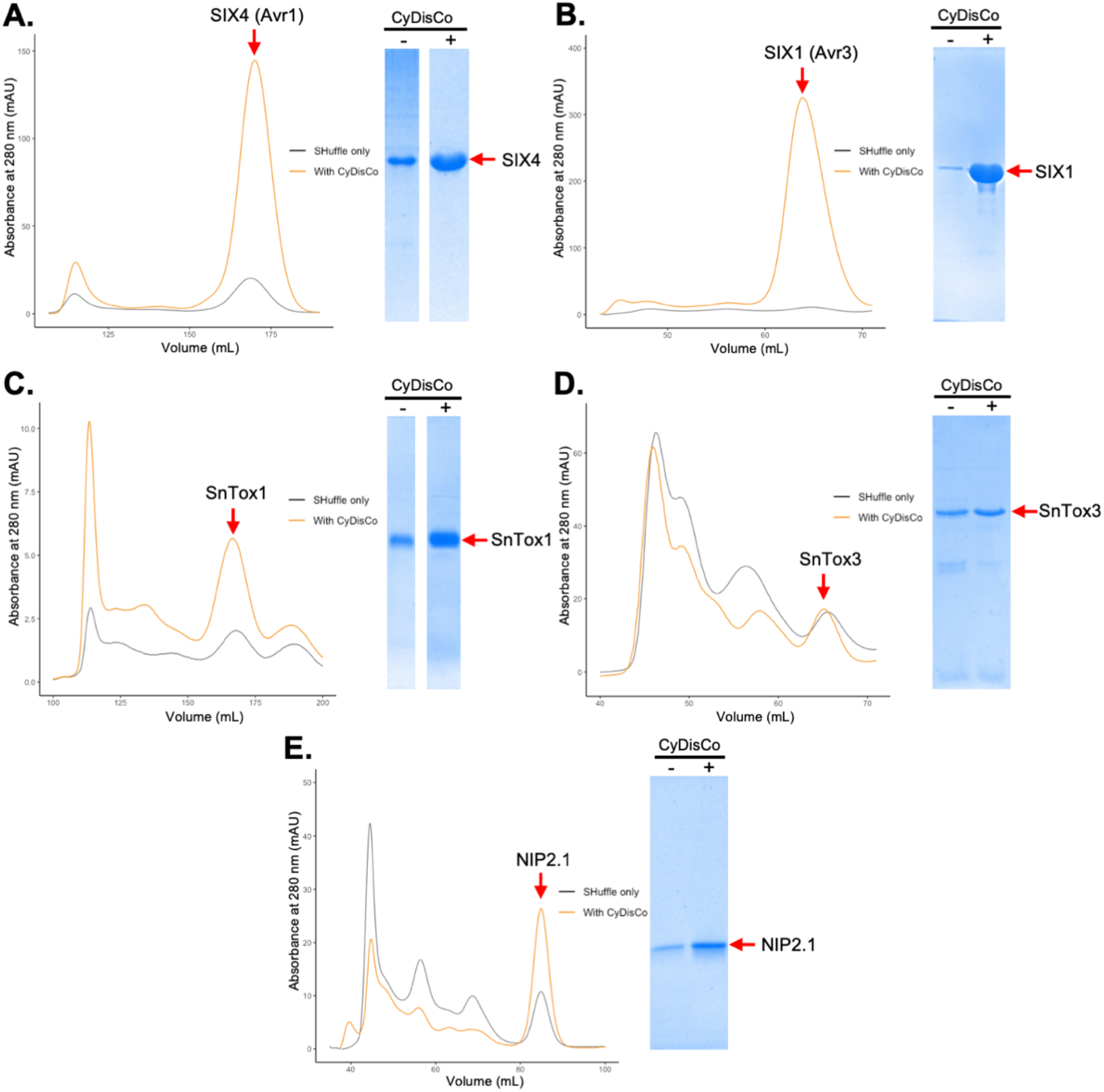
CyDisCo co-expression in SHuffle *E. coli* can be used effectively to produce various disulfide-bonded fungal effector proteins. Left panel: Size-exclusion chromatograms (SEC) of effectors produced in SHuffle *E. coli*, following cleavage by 3C protease to remove their N-terminal fusions, with CyDisCo co-expression (orange) or without (black). The red arrow indicates the peak corresponding to the effector of interest. Right panel: Coomassie-stained SDS-PAGE gel showing equal volume loading of effector of interest (indicated by red arrow) expressed with (+) or without (-) CyDisCo corresponding to the protein peak from SEC. Chosen effectors are **(A)** SIX4 (Avr1) and **(B)** SIX1 (Avr3) from *Fusarium oxysporum* f. sp. *lycopersici* and **(C)** SnTox1 and **(D)** SnTox3 from *Parastagonospora nodorum*, and **(E)** NIP2.1 from *Rhynchosporium commune*.

### A modified fungal-specific CyDisCo system for improved soluble expression of disulfide-rich fungal effectors

The Erv1p and human PDI pair of CyDisCo was previously reported to be the most effective at increasing protein yield (Gaciarz et al. 2016). However, this system has been used predominantly to enhance the production of disulfide-rich human proteins such as antibodies, human growth factor and perlecan (Matos et al. 2014; Gaciarz et al. 2016; Sohail et al. 2020). Recently, a modified CyDisCo system was utilised to produce disulfide-rich conotoxins from cone snails, whereby an additional conotoxin-specific PDI from *Conus geographus* was included with the CyDisCo components (Nielsen et al. 2019).

We have shown that CyDisCo benefits the production of numerous disulfide-rich fungal effectors. Despite this advance, the yield for some effectors, such as FovSIX6, remained low (0.2 mg/L) and we wanted to investigate whether the CyDisCo system could be modified and improved to benefit the production of recalcitrant disulfide-rich fungal effectors.

We substituted the human PDI with a PDI from *Fol*, as the amino acid sequence of human PDI is substantially divergent from fungal PDIs (Supplementary Fig. S3A). We also selected a sulfhydryl oxidase (Erv2) that localises in the endoplasmic reticulum (ER) of fungi to co-express with PDI in place of ERV1p. Erv2 is a fungal-specific membrane-bound sulfhydryl oxidase that catalyses disulfide bonds *de novo* within the ER (Sevier et al. 2001; Sevier and Kaiser 2006). When overproduced in yeast, Erv2 forms mixed disulfide bonds with yeast PDI, suggesting a transient association between the two proteins (Sevier et al. 2001). We selected Evr2 for two reasons, firstly, PDIs localise to the ER and would not interact with Erv1p-like sulfhydryl oxidases, which localise in the mitochondria (Lange et al. 2001; Ellgaard and Ruddock 2005). Secondly, the presence of signal peptides in fungal effectors indicate that they are trafficked through the ER and Golgi secretory pathway (Petre and Kamoun 2014).

BlastP searches of the *Fol* genome (Ma et al. 2010) using yeast PDI and Evr2 as queries, indicated that four putative PDI proteins and two Erv2-like proteins were present in *Fol* (Supplementary Fig. S3A and B). To select the most appropriate proteins for co-expression studies in *E. coli*, we made use of RNAseq data from *Fol* infections of tomato (Fig. 4A). This demonstrated that FOXG_00140 (FolPDI) and FOXG_09255 (FolErv2) were upregulated during infection, and these proteins were subsequently selected for expression trials (Fig. 4B).

**Fig. 4.**
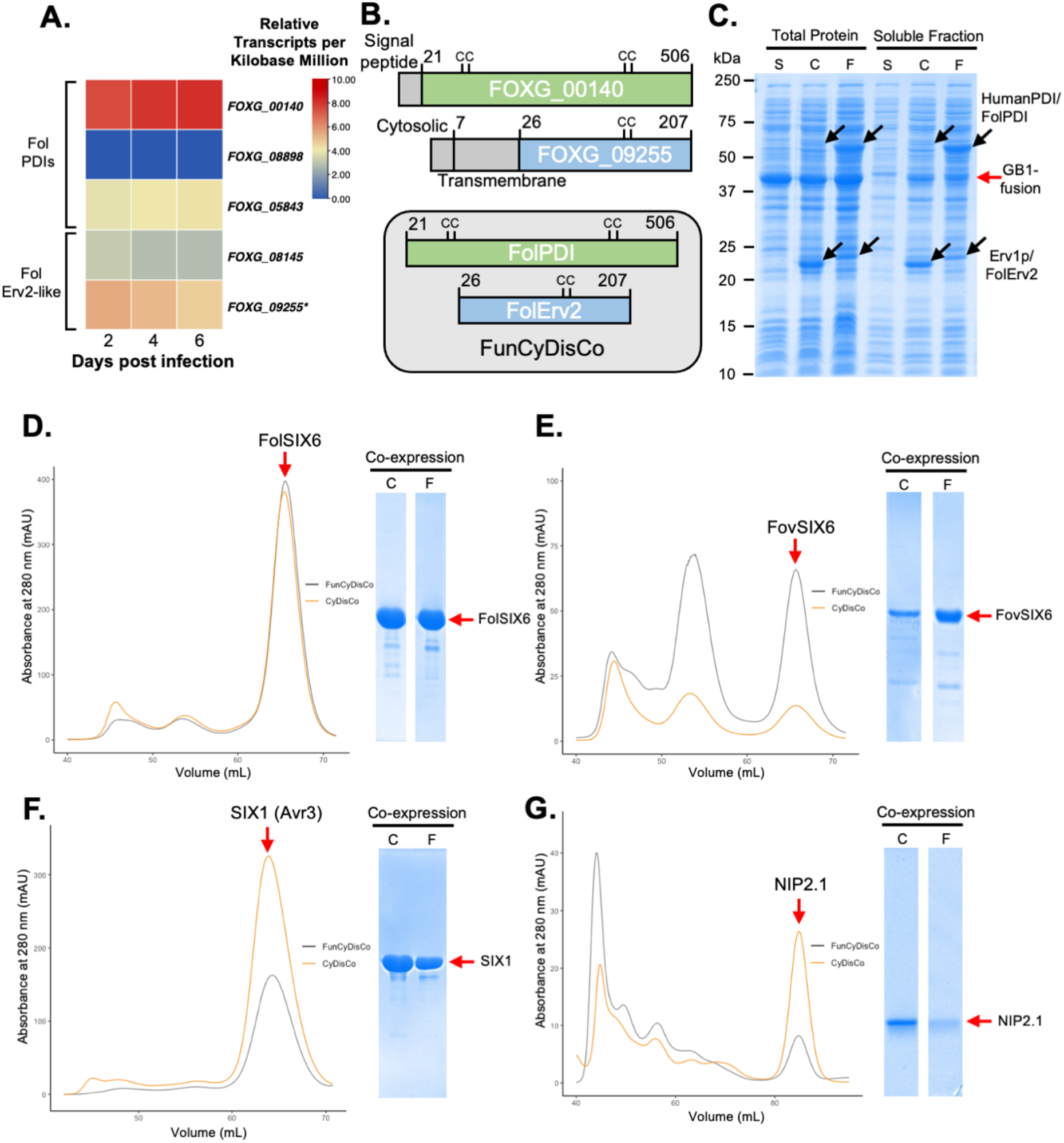
A fungal-specific CyDisCo system (FunCyDisCo) further improves yields of some effectors. **(A)** RNAseq analysis of *Fusarium oxysporum* f. sp. *lycopersici* protein disulfide isomerases (PDIs) and sulfhydryl oxidases (Erv2-like) during Fol infection of tomato. Transcripts of Fol PDIs (FOXG_00140, FOXG_08898, FOXG_05843) and sulfhydryl oxidases (FOXG_08145, FOXG_09255) measured at 2, 4 and 6 days post infection are shown. Relative scale represents Transcripts Per Kilobase Million (TPM) with values ranging from 0 to 281 TPM. **(B)** Schematic of selected components of FunCyDisCo (top panel) and domains that are expressed (bottom panel). **(C)** Representation of total protein and soluble fractions following expression of a GB1-fusion-effector of interest with the CyDisCo (C) or FunCyDisCo (F) co-expression systems, or SHuffle alone (S). Black arrows indicate overexpression of PDI or sulfhydryl oxidase components. Size-exclusion chromatogram and SDS-PAGE analysis of the *Fusarium oxysporum* effectors **(D)** FolSIX6, **(E)** FovSIX6, **(F)** SIX1(Avr3) and **(G)** *Rhynchosporium commune* effector NIP2.1 recombinant proteins co-expressed with CyDisCo (C) or FunCyDisCo (F). Red arrows indicate the protein of interest peak on the size-exclusion chromatogram and band on SDS-PAGE gel. *FOXG_09255 was incorrectly annotated. The FOXG_09255 sequence has been corrected based on RNASeq data and can be found in Supplementary Table S2.

To assess whether we could improve the CyDisCo system for production of disulfide-rich fungal effectors in *E. coli, F. oxysporum* effectors FolSIX6, FovSIX6 and SIX1 with an N-terminal 6xHisGB1 tag, and NIP2.1 from *R. commune* with an N-terminal 6xHis tag in SHuffle *E. coli* were co-expressed with either CyDisCo or the modified fungal-specific CyDisCo (FunCyDisCo) containing FolErv2 lacking the N-terminal transmembrane domain and FolPDI lacking the signal peptide (Fig. 4B), and purified them (side-by-side) using the same approaches detailed above. To confirm CyDisCo/FunCyDisCo components were expressed in a soluble form, total and clarified lysates were analysed by SDS-PAGE (Fig. 4C). For FolSIX6, the co-expression of CyDisCo or FunCyDisCo were equally effective, each resulting in a yield of ∼2 mg per litre of culture, a 5-fold increase compared to SHuffle alone (Fig. 4D). FovSIX6 co-expressed with FunCyDisCo resulted in a yield of ∼0.6 mg per litre of culture, a 3-fold improvement in yield compared to co-expression with CyDisCo and 15-fold improvement compared to SHuffle alone (Fig. 4E). For SIX1, co-expression with FunCyDisCo resulted in a 13-fold improvement in yield compared to SHuffle alone and a 2-fold decrease in protein yield when compared to CyDisCo (Fig. 4F). NIP2.1 co-expressed with FunCyDisCo resulted in a yield of ∼0.06 mg/L, which was similar to the yields obtained from SHuffle alone, but a 2.5-fold decrease compared to CyDisCo (Fig. 4G). Collectively, these results suggest the use of CyDisCo or FunCyDisCo co-expression systems can both improve yields of disulfide-rich effectors compared to SHuffle *E. coli* alone, however the choice of which system works best is protein specific.

### Co-expression of disulfide-rich fungal effectors in non-redox mutant *E. coli* strains

We have shown the CyDisCo and FunCyDisCo co-expression systems are effective at improving yields for numerous disulfide-rich fungal effectors produced in SHuffle *E. coli* compared to SHuffle alone. In a previous report, the CyDisCo system could be used to produce disulfide-bonded antibody fragments in different *E. coli* strains (Gaciarz et al. 2016). This could allow greater flexibility in the choice of *E. coli* background for the expression of disulfide-rich effectors, which might be advantageous for different applications.

We therefore investigated if improvements to the yield of disulfide-rich fungal effectors can be made using the CyDisCo and FunCyDisCo systems expressed in non-redox mutant strains such as BL21(DE3). FolSIX6 lacking the signal peptide with an N-terminal 6xHisGB1 tag was expressed in BL21(DE3) by itself, or co-expressed with CyDisCo or FunCyDisCo systems, and was purified simultaneously from both using nickel affinity chromatography. However, we were unable to produce high quantities of FolSIX6 with either co-expression system in BL21(DE3). We were also unable to confirm the soluble production of CyDisCo/FunCyDisCo components (Supplementary Fig. S4). Collectively, in our hands, the CyDisCo and modified FunCyDisCo systems were not transferable into BL21(DE3) *E. coli*.

### Recombinant SIX4 (Avr1) causes cell death in *I*-containing tomato cultivars

The adoption of the CyDisCo/FunCyDisCo system has facilitated the structural elucidation of numerous fungal effectors in our lab (structures to be presented elsewhere). Here, however, we present data to show that these high-quality/purity proteins have applications outside of structural studies. Previously, we used a protein-mediated phenotyping approach to study the necrotrophic effector SnTox1 and SnTox3 in wheat (Zhang et al. 2017; Outram et al. 2021; Sung et al. 2021). Here, we were interested in determining whether the effectors produced using our enhanced production system could be used to study effector recognition. We demonstrated that purified SIX4 (Avr1) protein infiltrated into cotyledons caused cell death in a tomato cultivar that contained the *I-*resistance gene (M82). Importantly, cell death was not observed when the same protein was infiltrated into a tomato cultivar lacking *I* (Moneymaker) (Fig. 5). This demonstrates the capacity for *E. coli*-produced SIX4 (Avr1) to be recognised by the I resistance protein in the native tomato system.

**Fig. 5.**
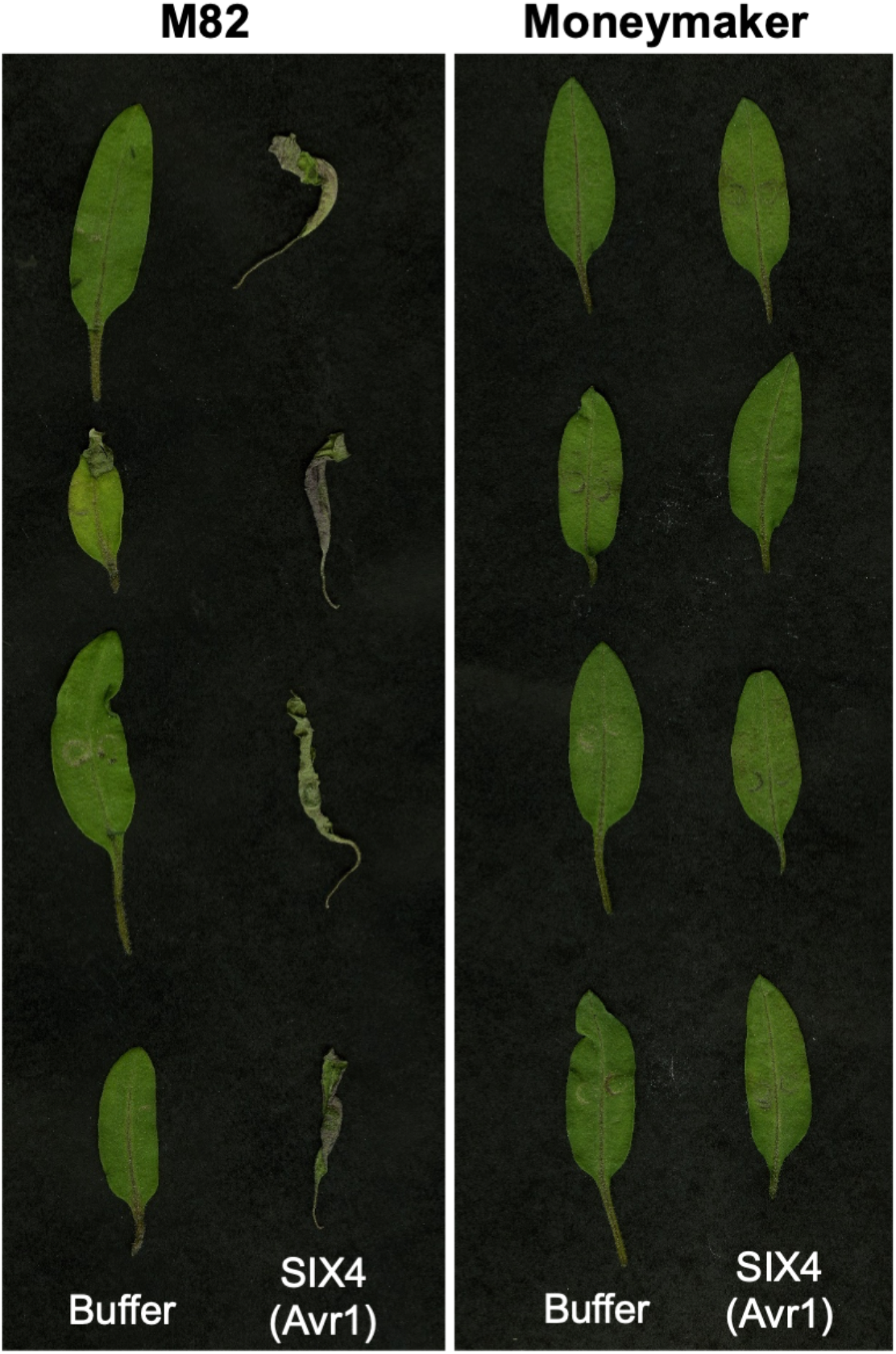
*Escherichia coli*-produced SIX4 (Avr1) causes cell death when infiltrated into tomato cultivars containing the resistance gene *I*. SIX4 (Avr1) (at a concentration of 100 μg/mL) and a buffer control were syringe-infiltrated into 10-day old tomato cotyledons from cultivars M82 (containing *I*) and Moneymaker (lacking *I*). Cotyledons were harvested and imaged 4 days post infiltration.

## Discussion

Here, we demonstrate that the CyDisCo co-expression strategy has the capacity to significantly increase the yield of functional disulfide-rich effectors when produced in SHuffle *E. coli*. Of the eight effectors we trialled, all could be expressed and purified using CyDisCo and seven displaying improved yields and purity compared to SHuffle alone. Our tailored FunCyDisCo outperformed SHuffle alone for the three *F. oxysporum* effectors studied, but showed effector-specific differences compared to CyDisCo.

The basis of the CyDisCo co-expression approach is to mimic (albeit loosely) eukaryotic secretory pathways within a prokaryotic host. PDI and sulfhydryl oxidases proteins are known to function together to assist protein folding through disulfide-bond formation and correct pairing of disulfide bonds (Sevier 2010). There is some evidence that these proteins are also important in pathogenic fungi. For example, PDI1 from *Ustilago maydis* is crucial for the correct folding of a pool of secreted disulfide-rich proteins important for virulence (Marin-Menguiano et al. 2019). We attempted to further tailor this system with the introduction of FunCyDisCo co-expression, using *Fol* PDI and Erv2 isoforms identified in RNAseq data from *Fol-*infected tomato. Our data for FunCyDisCo showed mixed success compared to CyDisCo. One potential reason for this variation is isoform specificity. In *Saccharomyces cerevisiae*, there are more than five PDI-like proteins and two sulfhydryl oxidases localised to the endoplasmic reticulum, each preferentially aiding the disulfide-bond formation of different proteins (Frand and Kaiser 1999; Norgaard et al. 2001; Sevier and Kaiser 2006). In *Fol*, four PDI and two Erv2-like homologs can be identified. While RNAseq data from host infection were used to guide our selection, it is plausible other homologs or combinations could result in better effector yields.

Further improvements may also be derived from the introduction of additional accessory proteins and chaperones that are not specifically involved in disulfide bond formation. For example, Lhs1, an HSP70 chaperone from *Magnaporthe oryzae* is crucial for the proper processing of secreted proteins, with *Lhs1* knockouts exhibiting lower levels of secreted effector proteins and severely reduced pathogenicity (Yi et al. 2009). Other systems involving co-expression of accessory proteins and chaperones have been successfully utilised to express complex proteins in *E. coli*, such as RuBisCo. The simultaneous co-expression of a plant chaperonin and four assembly factors has been reported to produce ∼12-fold higher yields of functional RuBisCo (Wilson et al. 2019). The incorporation of general accessory proteins and chaperones that aid the oxidative pathway for disulfide-bond formation may tailor the co-expression system for a given protein. However, due to the complexity of different oxidation pathways, pinpointing which proteins to co-express with a given disulfide-rich effector is difficult.

Eukaryotic proteins produced in a prokaryotic system often end up in inclusion bodies due to the lack of folding machinery and a rapid rate of protein synthesis preventing correct protein folding (Widmann and Christen 2000). Incorporation of eukaryotic components for co-expression in *E. coli* to mimic eukaryotic disulfide-bond formation raises the question: Why not use a eukaryotic system directly? Eukaryotic expression systems like yeast, have been used successfully to express soluble disulfide-rich effectors in quantities necessary for structural characterisation, such as AvrLm4–7 from *Leptosphaeria maculans* and Ecp6 from *Cladosporium fulvum* (Sanchez-Vallet et al. 2013; Blondeau et al. 2015). However, some disulfide-rich proteins were produced in lower quantities when expressed in yeast compared to *E. coli*, such as SnTox1, SnTox3 and ToxB (Liu et al. 2009; Liu et al. 2012; See et al. 2019; Outram et al. 2021). This demonstrates that choosing expression systems for producing disulfide-rich effectors is not a ‘one size fits all’ approach and multiple expression systems and strategies should be considered in the early stages of recombinant effector protein production.

With prokaryotic expression systems being cheap and accessible to many laboratories, we believe our combined strategy of SHuffle *E. coli* strain, GB1 solubility tag and CyDisCo or FunCyDisCo co-expression systems would assist the characterisation of disulfide-rich effectors from a broad range of plant pathogens.

## Materials and Methods

### Vectors and gene constructs

Fungal effector gene DNA sequences were codon optimised for expression in *E. coli* and synthesised by Integrated DNA Technologies (IDT, Coralville, Iowa, USA) (Supplementary Table S2). All genes were cloned into the modified, Golden Gate-compatible, pOPIN expression vector (Bentham et al. 2021). The final expression constructs contained either a N-terminal 6xHis-tag or 6xHis-GB1-tag followed by a 3C protease recognition site. The Golden Gate digestion/ligation reactions and cycling were carried out as described by Iverson et al. (2016).

DNA sequences that encode the yeast Erv1p and human PDI (CyDisCo) and *Fol* Erv2 *Fol* PDI (FunCyDisCo) were codon optimised using the tool provided by IDT, and synthesised by Twist Bioscience (San Francisco, California, USA) (Supplementary Table S2). The Yeast Erv1p and Human PDI pair, and *Fol* Erv2 and *Fol* PDI pair were cloned into a modified Golden Gate compatible pACYC184 vector from the EcoFlex Kit (Moore et al. 2016), which was a gift from Paul Freemont (Addgene kit #1000000080). The Golden Gate digestion/ligation reactions and cycling were carried out as described by the kit (Moore et al. 2016). All plasmid constructs were sequence verified by sequencing.

### Protein expression and purification

Sequence verified effector constructs (∼100 ng plasmid DNA) were chemically transformed into SHuffle T7 Express C3029 (New England Biolabs (NEB), Ipswich, Massachusetts, USA) or BL21(DE3) C2527 competent *E. coli* (NEB) using the heat shock protocol provided by the manufacturer and the transformants grown on LB agar plates supplemented with ampicillin (100 µg/mL) at 37°C for 16 h. For CyDisCo/FunCyDisCo co-expression, the effector of interest and CyDisCo/FunCyDisCo constructs (∼100 ng plasmid DNA) were transformed simultaneously and the transformants grown on LB agar plates supplemented with ampicillin (100 µg/mL) and chloramphenicol (35 µg/mL) at 37°C for 16 h. Colonies were used to inoculate Luria-Bertani (LB) media supplemented with required antibiotics and grown overnight at 37°C (BL21(DE3)) or 30°C (SHuffle) with shaking at 220 rpm. These small-scale overnight cultures were used to inoculate 1 L of Teriffic Broth media (24 g/L yeast extract, 12 g/L tryptone, 0.5% (v/v) glycerol, 0.017 M KH_2_PO_4_, 0.072 M K_2_HPO_4_) in a 2 L baffled flask supplemented with required antibiotics and 200 µL of Antifoam 204 (Sigma-Aldrich Inc., St. Louis, Missouri, USA). Large-scale cultures were incubated at 37°C (BL21(DE3)) or 30°C (SHuffle) with shaking at 220 rpm. Cultures were induced with a final concentration of 200 µM isopropyl β-D-1-thiogalactopyranoside (IPTG) once an OD_600_ of 0.6 was reached and incubated at 16°C with shaking at 220 rpm for a further 16 h. Cells were harvested by centrifugation at 4000 xg for 10 min at 4°C. Pellets were resuspended in 50 mM HEPES pH 8.0, 300 mM NaCl, 10% (v/v) glycerol, 1 mM phenylmethylsulfonyl fluoride (PMSF) and lysed by sonication using an amplitude of 40% (10 seconds on, 20 seconds off). The lysed cells were centrifuged at 20000 xg for 40 min to clarify the lysate. The protein of interest was purified further by immobilised metal affinity chromatography (IMAC) using a 5 mL HisTrap FF crude nickel column (Cytiva, Marlborough, Massachusetts, USA). The column was washed using a buffer containing 50 mM HEPES pH 8.0, 300 mM NaCl, 30 mM imidazole, prior to elution using either gradient elution or isocratic elution (dependent on the effector protein) with a buffer containing 50 mM HEPES pH 8.0, 300 mM NaCl, and 250 mM imidazole. Eluted fractions were analysed by SDS-PAGE and fractions containing the protein of interest were dialysed to remove imidazole and incubated with 6xHis-tagged 3C protease (150 µg) overnight at 4°C to cleave off the N-terminal fusion from the effector proteins. Cleavage was confirmed via SDS-PAGE, and the cleaved protein of interest was separated from the N-terminal fusion tag, any uncleaved protein and 6xHis-tagged 3C protease using IMAC, and subsequently purified further by size-exclusion chromatography (SEC) using either a HiLoad 16/600 or HiLoad 26/600 Superdex 75 pg column (GE Healthcare) equilibrated with a buffer containing 10 mM HEPES pH 8.0 and 150 mM NaCl. Proteins were concentrated using a 3 or 10 kDa molecular weight cut-off Amicon centrifugal concentrator (MilliporeSigma, Burlington, Massachusetts, USA), snap-frozen in liquid nitrogen and stored at -80°C for future use.

### Intact protein mass spectrometry (MS)

Proteins were adjusted to 10 µM in 0.1% (v/v) formic acid (FA) for HPLC-MS analysis. The samples were then injected onto an Agilent UHPLC system. Each sample was first desalted for 2 min on an Agilent (Santa Clara, California, USA) C3 trap column (ZORBAX StableBond C3) at a flow rate of 500 µL/min at 95% the buffer A (0.1% FA, v/v) and 5% the buffer B (0.1% FA and 100% ACN) followed by separation over 8 min using a 5–80% (v/v) gradient of buffer B at a flow rate of 500 µL/min. Eluted material was analysed using a Orbitrap Fusion™ Tribrid™ mass spectrometer (Thermo Fisher Scientific, Waltham, Massachusetts, USA). MS acquisition was performed using the Intact Protein Mode script. The acquisition was performed across m/z 200-4000 with an accumulation time of 1 s. Data were analysed using the Free Style v.1.4 (Thermo Fisher Scientific) protein reconstruct tool across a mass range of m/z 500 – 2000. The expected sizes of the proteins were searched.

### Circular dichroism (CD) spectroscopy

The CD spectra of purified effectors of interest were recorded on a Chirascan spectrometer (Applied Photophysics Ltd., UK) at 20ºC. Samples were diluted to 10 µM in a 20 mM sodium phosphate buffer at pH 8.0. Measurements were taken at 0.5 nm wavelength increments from 190 nm or 200 nm to 260 nm at a scanning speed of 50 nm/min. A cell with a pathlength of 1 mm, a bandwidth of 0.5 nm and response time of 4 s were used, with 3 accumulations. The data were averaged and corrected for buffer baseline contribution, and visualised using the webserver CAPITO tool with data smoothing (Wiedemann et al. 2013).

### Tomato infiltration assays

Tomato seeds were sown in seed raising mix and grown in a controlled environment chamber with a 16-h day/8-h night cycle at 22°C. Purified SIX4 (Avr1) protein was diluted in water to 0.1 mg/mL. Syringe infiltrations of the cotyledons of 10-d old tomato seedlings were conducted with 100 µl of protein or buffer (10 mM HEPES pH 8, 150 mM NaCl diluted 1/100). Cotyledons were harvested and imaged at 4 days post-infiltration (dpi).

## Acknowledgements

This work was supported by the Australian Research Council (ARC; DP180102355 P.S.; DP200100388 D.J./S.W.) and the Australian Academy of Science (Thomas Davies Grant). S.W. was funded by an ARC Future Fellowship (FT200100135) and is supported by the ANU Future Scheme (35665). L.M. was funded by an ARC Discovery Early Career Researcher Award (DE170101165). E.C. and A.S. were a recipient of the AINSE Honours Scholarship Program and D.Y. holds an AINSE Postgraduate Research Award. The vectors containing CyDisCo and FunCyDisCo will be deposited with Addgene for easy access by the research community. The mass spectrometry analysis was carried out at the joint mass spectrometry facility at The Australian National University. We thank Adam J. Carrol and Joseph Boileau for their technical assistance with the mass spectrometry experiments.

