## Supplemental for "Optimised production of disulfide-bonded fungal effectors in *E. coli* using CyDisCo and FunCyDisCo co-expression approaches"

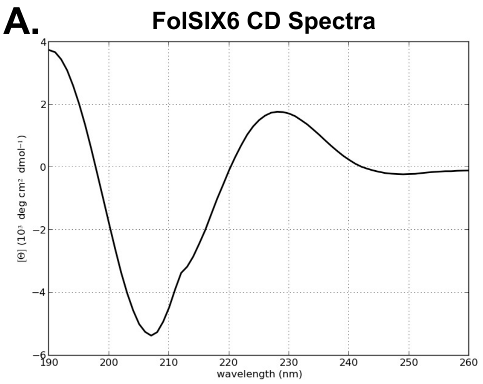


Secondary structure elements based on CD spectra

| helix | β-strand | irregular |
|---|---|---|
| 0.00 | 0.38 | 0.54 |

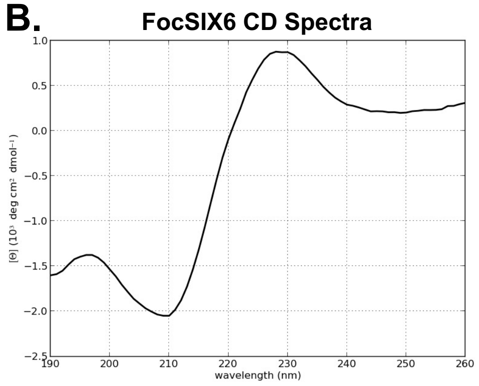


Secondary structure elements based on CD spectra

| helix | β-strand | irregular |
|---|---|---|
| 0.00 | 0.49 | 0.56 |

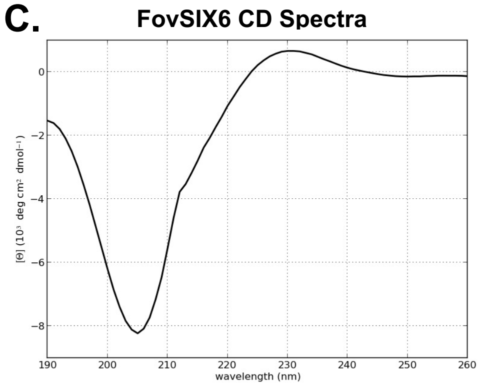


Secondary structure elements based on CD spectra

| helix | β-strand | irregular |
|---|---|---|
| 0.01 | 0.33 | 0.61 |

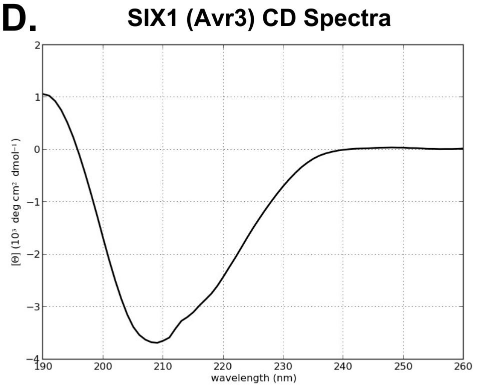


Secondary structure elements based on CD spectra

| helix | β-strand | irregular |
|---|---|---|
| 0.02 | 0.43 | 0.56 |

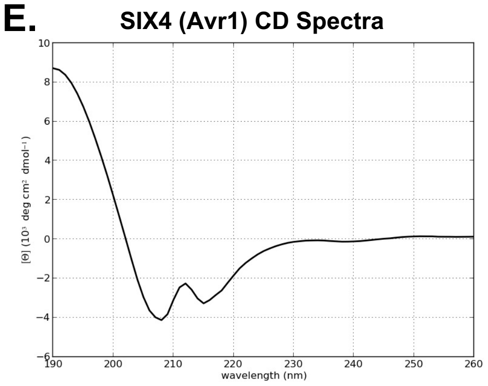


Secondary structure elements based on CD spectra

| helix | β-strand | irregular |
|---|---|---|
| 0.04 | 0.44 | 0.50 |

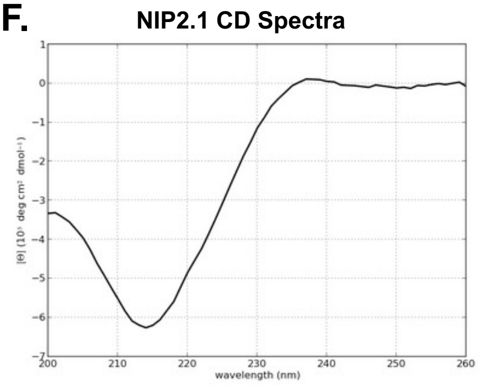


Secondary structure elements based on CD spectra

| helix | β-strand | irregular |
|---|---|---|
| 0.26 | 0.38 | 0.36 |

**Supplementary Fig. S1. Circular Dichroism (CD) spectra of disulfide-rich fungal effectors produced by SHuffle with CyDisCo co-expression.** CD spectra of the *Fusarium oxysporum* f. sp. *lycopersici* effectors **(A)** FolSIX6, **(B)** FocSIX6, **(C)** FovSIX6, **(D)** SIX1 (Avr3), **(E)** SIX4 (Avr1) and **(F)** the *Rhynchosporium commune* effector NIP2.1 recombinant proteins plotted, and secondary structure elements analysed using the CAPITO webserver (Wiedemann et al. 2013).

**A. FolSIX6**

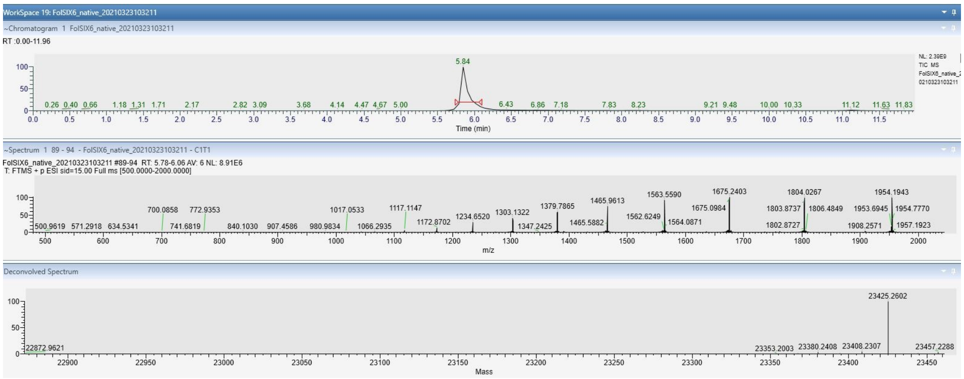


**B. FocSIX6ΔPD**

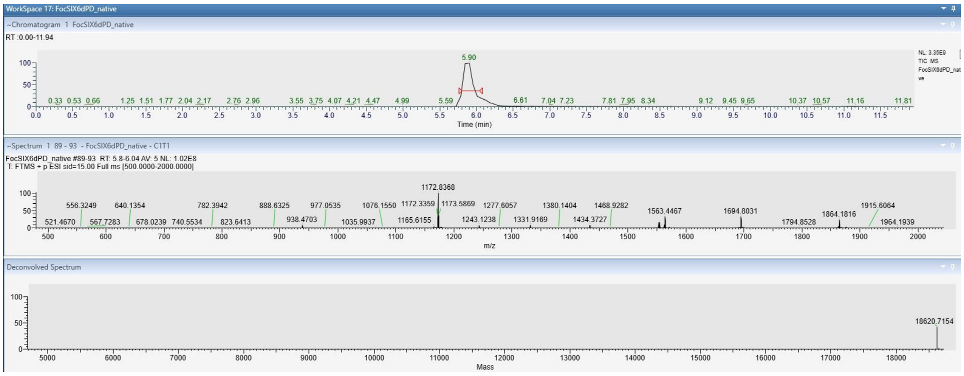


**C. FovSIX6**

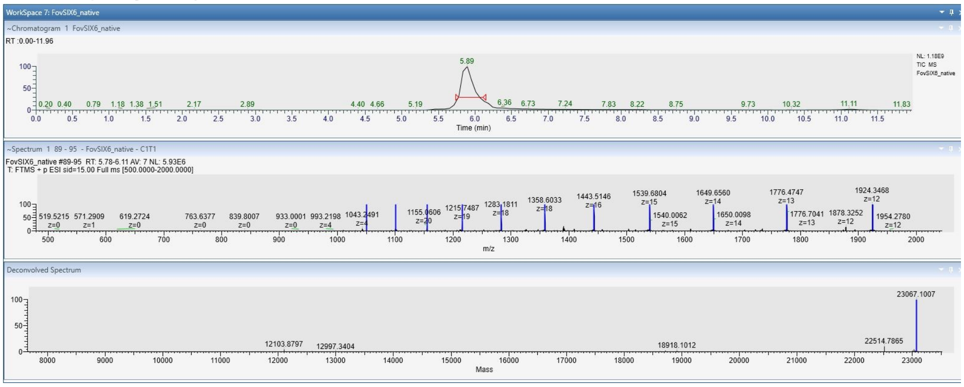


**D. SIX1 (Avr3)**

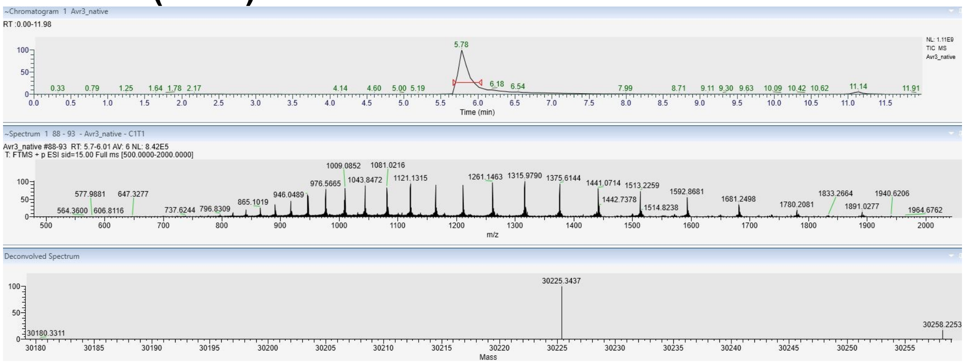

### E. SIX4 (Avr1)

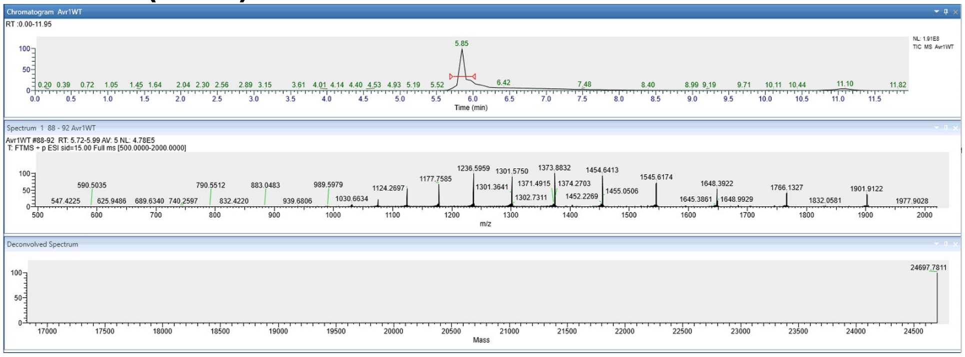


### F. NIP2.1

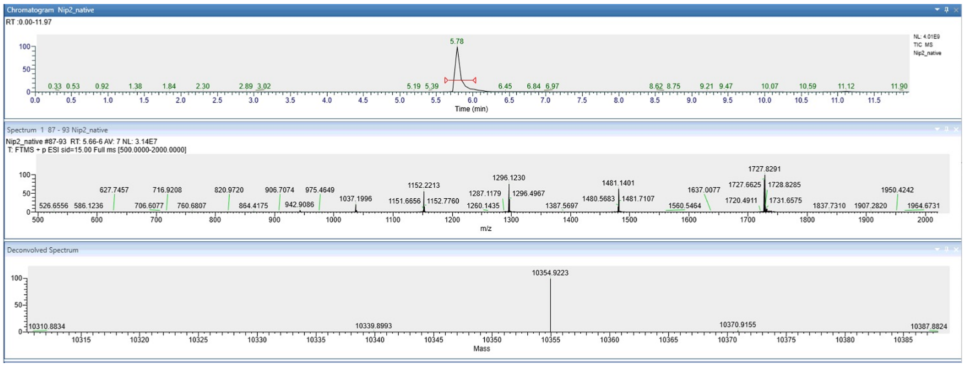


**Supplementary Fig. S2. Native intact protein mass spectrometry analysis to determine oxidation state of recombinant proteins.** Native intact protein mass spectrometry analysis of **(A)** FolSIX6, (B) FocSIX6ΔPD, (C) FovSIX6, (D) SIX1 (Avr3), (E) SIX4 (Avr1) and (F) NIP2.1 produced SHuffle with CyDisCo co-expression. Comparison with theoretical masses when cysteines are oxidised or reduced can be found in Supplementary Table S1. Intact mass spectrometry on SnTox1 is reported by Zhang et al., 2017, and for SnTox3 by Zhang et al., 2017 and Outram et al., 2021.

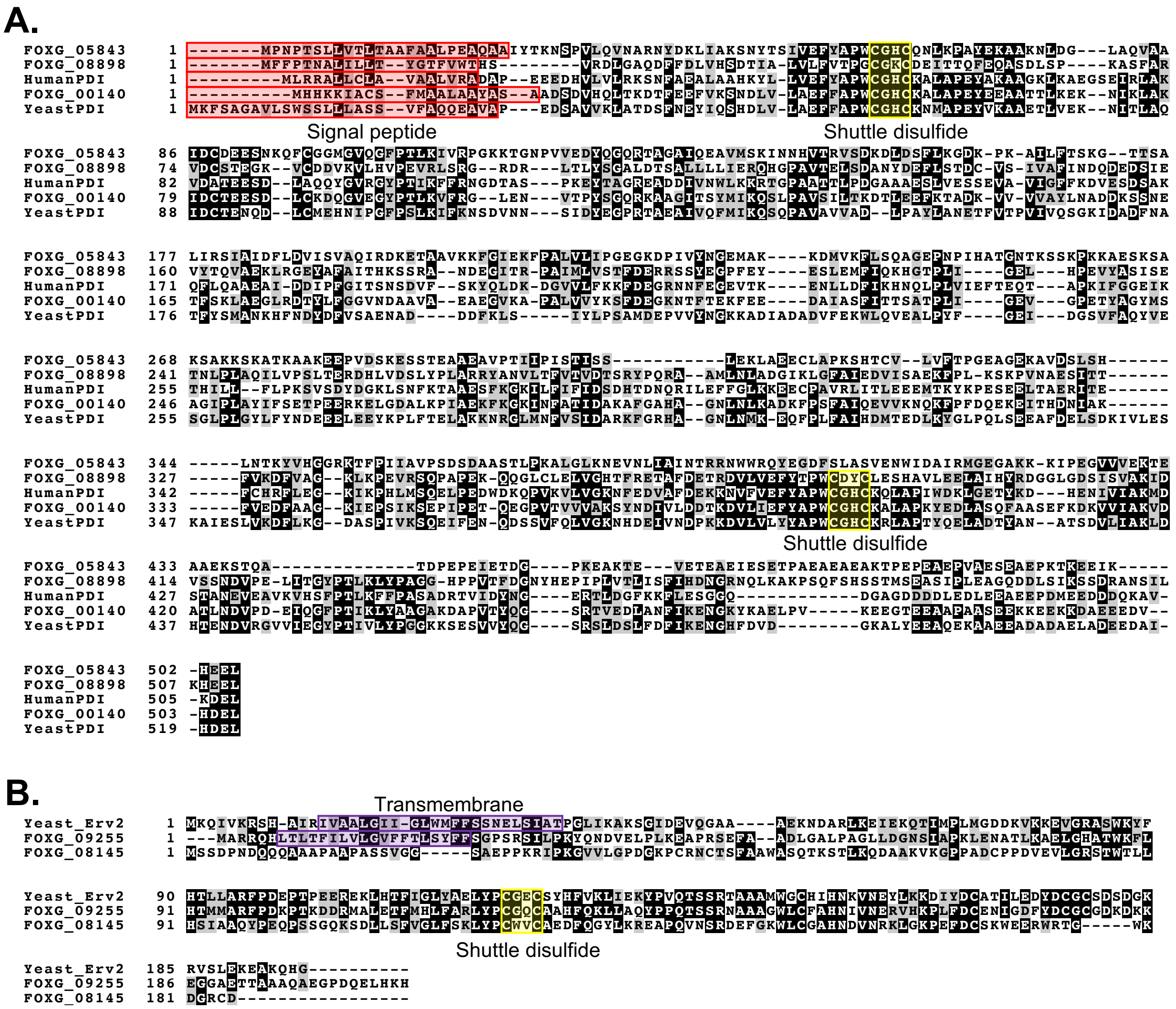


**Supplementary Fig. S3. Sequence alignment of SIX6, PDI and Erv2-like proteins. (A)** Alignment of Human and Yeast PDIs, and PDI-like proteins from Fol. **(B)** Alignment of Erv2 and Erv2-like proteins from Fol. Signal peptides are shown in red, transmembrane domains are shown in purple, and redox active disulfide domains (Shuttle disulfides) are highlighted in yellow.

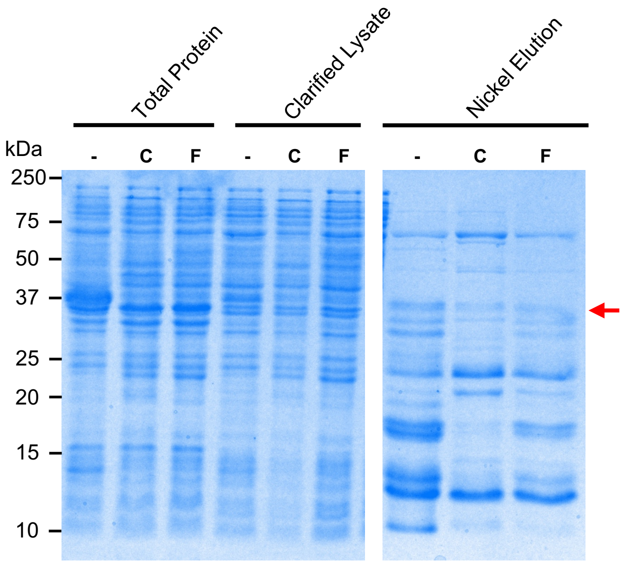


**Supplementary Fig. S4. FolSIX6 cannot be produced in high yields in BL21(DE3) *E. coli* despite the use of CyDisCo or FunCyDisCo co-expression systems.** Coomassie-stained SDS-PAGE gel showing total and soluble (Clarified lysate) proteins, and proteins captured by immobilised metal affinity chromatography (IMAC) from BL21(DE3) *E. coli* expressing 6xHisGB1-FolSIX6 without any co-expression system (-) or co-expressed with CyDisCo (C) or FunCyDisCo (F). The red arrow points to the 6xHisGB1-FolSIX6 protein band.

**Supplementary Table S1. Fungal effectors chosen in this study and masses determined by mass spectrometry**

| Effector | Fungal Pathogen | Protein length (amino acids) | Cysteines | Determined Mass (Da) | Predicted Reduced Mass (Da) | Predicted Oxidised Mass (Da) |
|---|---|---|---|---|---|---|
| FolSIX6 | *Fusarium oxysporum* f. sp. *lycopersici* | 209 | 8 | **23425.3**^ | 23433.3 | **23425.3** |
| FocSIX6 (ΔPD) | *Fusarium oxysporum* f. sp. *cubense* | 211 (168) | 8 (8) | - **18620.7** | - 18628.8 | - **18620.8** |
| FovSIX6 | *Fusarium oxysporum* f. sp. *vasinfectum* | 208 | 8 | **23067.1**^ | 23075.1 | **23067.1** |
| SIX1 (Avr3) | *Fusarium oxysporum* f. sp. *lycopersici* | 263 | 8 | **30225.3**^ | 30233.3 | **30225.3** |
| SIX4 (Avr1) | *Fusarium oxysporum* f. sp. *lycopersici* | 225 | 6 | **24698.8**^ | 24703.8 | **24697.8** |
| SnTox1 | *Parastagonospora nodorum* | 100 | 16 | -* | - | - |
| SnTox3 | *Parastagonospora nodorum* | 210 | 6 | -* | - | - |
| NIP2.1 | *Rhynchosporium commune* | 93 | 6 | **10354.9**^ | 10361.0 | **10355.0** |

*Mass spectrometry reported for SnTox1 by Zhang et al. (2017), and for SnTox3 by Zhang et al. (2017) and Outram et al. (2021).

^Proteins contain additional GPM residues on the N-terminus and S residue on the C-terminus as a result of 3C protease cleavage and cloning scars.

**Supplementary Table S2. Genes used in this study**

| Name | Sequence |
|---|---|
| FolSIX6 | TAGGTCTCCAATGGGTCCCTTAGCCCAAACAGAATCCGAGTCGGCAGACGTCGCTGAACATAC<br>AATCAATTATATCGACATTGCCCCTGAAGAATTTGAACCGCCCAAAGCTAATTTGTCATCTCTG<br>GTGAGTCGTGACACGCTTCCTGTCAGTACCTGTCCTGCGGGTCAGAAATACGATCGTTCCGTGT<br>GTTACAAGGCAGACAAAATTCGTAGCTTTTGTGTCGCAAACCCTCGTAGCAACCGTGAGAAGA<br>TTACCGACACACCGTGTCAGCCCCGTGAAATCTGTGTGCAACGCAATCTTTCCAACGGCAAGA<br>GTTTCGCTAAGTGTATCCCCATTGTAGACCTGGTGGAATGGAAGACATCCGCAAATGGGAATA<br>AAGAAGGCTGTACTACAACGTCCGTGAATCCGGCTGGGTACCATCACCTTGGTACTATTGTTTA<br>CGATATCAATAAGAATCCTATCGAAGTTGATAAAATCTCGTACTTCGGCGAGCCGGGAAATGT<br>AAACGAGGGCATTGGTGGCAGCACAAGCTATTTTAGTAGTGACAACTTTCAATTTTCTAAGTCC<br>CGCTACATGAAAACTTGTATTTTCAGTGGTGGGTACGGGAATCTTAACGCCTATACGTGGAGCT<br>GGGAATCTTGGAGACCGT |
| FocSIX6Thrombin | TAGGTCTCCAATGTCTCCCTTGGATCCAGCCAAGACGCCCACGAGCCCGCCAGGAGCAGAACA<br>TACTCTGAACTACGTTGATATCACCCCTACCGGCCCAGAATTTGGCAATGTAAATGGCTCCTCG<br>GCGTTAGTACCTCGCAGTACACTGCCGCATACTGCGTGTCCAGCCGGGCAAACGTACGACCGC<br>TCAGTTTGCTACAACTCGCACACTATCCGTTCATTTTGCGTCGCTAATCCGCGTAGTAACCGTGA<br>ACAGATTACTGACACGCCATGTAACTCAGGAGAAGTATGTGTGCAGCGTCGCTTATCTAGTGG<br>TAAAAGCTACGCAAAATGTTTACCTGTCCGCGACTTAGTGTCGTGGCGCACGGACCCTGACGG<br>TGATAAGGAGGGCTGTACTACGGTTGAGGCGAACCCTATCGGGTATCACTCATTAGCGACGAT<br>GATCTATGATATCAACAATAACCCAATCCAAGTCGACAAGATCCGCTACTTAGGCGAGCCCGG<br>CGACGCTAACGAAGGTATTGGAGGAAGCGTTTCGAACTTCTCATCAGACCGCTTTCGTTTCACA<br>GGCTCTAATTACATGAAGGTGTGTGTATTTAGCGGTGGGTATGGAAATTTGAACGCCTACACC<br>TGGGTGTGGAACTCTTGGAGACCGT |
| FovSIX6 | TAGGTCTCCAATGGGTCCGTTAGCTCAAACCGAGTCGGAATCGGCGGATGTAGCTGAGCATAC<br>CATCAATTACATTGACATTGCGCCCGAGGAGTTTGAACCACCTAAGGCAAACTTGTCGTCATTA<br>GTGAGTCGCGACACATTACCTCCCACCTGCCCGCGCTACACGACATACGATGAGTCCATTTGTA<br>TCACGGCAGGCGTAGTACGCTCTTCTTGCGTTTTCACTGCAGATCCAGGAGTACGTCGTGGGG<br>TCAACGCGAATTGTAATAAGAACGAGATTTGCGTTGAACGCAACTTGTCTAACGGTAAACGCT<br>ATGCGAAGTGTATTCCCATCGCGCACTTGGTGGAATGGAAAACAAGTGCCGATGGCAACAAG<br>GAGGGCTGCACGACGACGAGCGTGAATCCAGCCGGTAATCATCATTTAGGCACGATCGTTTAC<br>GATGTTAACAAGAACCCGATTCAGGTTGATAAAATTTCATACTTTGGAGAACCCGGAAATGTG<br>GACGAGGGAATTGGTGGGTCTACCTCATACTTTTCTTCTAATTTGTTCCACTTCAGCAAGTCCCG<br>TTACATGAAAACGTGTATTTTCTCTGGAGGTTACGGGAACTTAAACGCTTATACGTGGTCCTGG<br>GTTTCTTGGAGACCGT |
| Avr1Thrombin | TAGGTCTCCAATGTTGCCTAAAGGAGAGGAGGGTGACATTATTGGTACTTTCAATTTCTCGTCC<br>AGCGACAGCCAACCCCTTAAAATCCACTGGGTCGATACGCCGGACTCATCTGGGAGCAATCTT<br>GTTCCCCGTTCCGCTCACACGGAGAGTGTATGCGTTCACGCCGGGACCGCTACAGGTGCTGAT<br>CTGCATTGGTTGAATGCGATCTGCACCGGGAAGTCTACATACACAGTGAATTGCGCCCCGGCA<br>GGCAACAAGAATGCTGGGTCTACGCACACAGGAACATGTCCGGCAGGTCAGGACTGTTTCCAA<br>TTAGAGCAGGTCGGAAACTTTTGGGGGGACCGTGAGCCAGATGCTACCTGTAGCCCGTCCAAT<br>ACGGTATTTGACGCCGTAGATGACAAGGAAGCTACGCATGTAAACGGCAAAGTTGTTACACGC<br>GCGGGGAAGCCGGGCATTGGGCGCAAGCTTATTCGTCTTAAGGCTCAGGTCTATCGTCGTGAT<br>GGTCACTATGGTCAGACCTCGCGCATGGGATTCTTTCGTAACGGCAAAGAGGTTTACCATATC<br>GACAACGTTGCCTCGATGGAACCCACTTGGAATTTTGACCCATCGAGTGACCAATCCTTTAGCT<br>TCTTTTTCACACCGGGACCCAACGCTTTCCGTATTCAAGGAACGCTTAATCTGGCCTCTTGGAG<br>ACCGT |

**Supplementary Table S2 Continued. Genes used in this study**

| Name | Sequence |
|---|---|
| Avr3Thrombin | TAGGTCTCCAATGCAGGAGGCGGCAGTCCGCGAGCCACAAATCTTTTTTAATCTGACTTACACC<br>GAGTATCTGGATAAAGTTGCCGCCTCACACGGCTCCCCGCCCGATAAAAGCGACTTGCCCTGG<br>AACGACACCATGGGGAGTTTCCCAGGTAATGAAACAGACGACGGGGTGCAGACTGAAACTGG<br>TTCGTCATTATCTCGCCGTGGGCATATTGTCAATCTGGTGCCGCGTTCCCCGTTCGGGGAGGAG<br>TCGCGCAATGATCGTGTGACGCAAGACATGTTACAGGCACTGCATGACCTGTGCGTCGAGCGT<br>TTCGGCACGGGGTATCGTGCGGTGAGCGGGTTGTGTTACACCGATCGTCGTGCTACACGTAAA<br>ATCGAATGTAATAAACCCTCTGTGCGCGAACGCGATCGCTCAGTAACCCGCGCGTGCCCGGAG<br>GGGCAGGAGTGCACCACCTTCAACGCCTACAATTTTCGTAACCGTCACCACCAGGTAACTTTCC<br>CCGTCTGCGGTCCTCGTATCGAGGTTAAGGATCGCCATGACATTGGGATTCATACTGAATGGC<br>AAGGAACGTGGTATCCGGAATCACCTAAATCACCAGGGACATACGATTATTTCGCCCAGATGG<br>CTGGGACGTTGAACGGTTACTTCGGTTACGATGGAGTATATAGCGATGGCTATAAAACTAGCT<br>CCCACGGCTACGGTCACAGTTGGTCATGTATTAATTGCCCCCGTGGGAAGGTGACAATTACGA<br>ACACGTATCGCGCTACATGGGCGTTCGGATACACTTCACCCCATTCTTGGAGACCGT |
| SnTox1 | TAGGTCTCCAATGAATGAAGGTATTCTGACATTTGAAGGTCTGGGTCTGGCACGTCGTCAGAC<br>CATTTGTCATACACCGGGTGGTAGCGGTTGTCGTGCAAGCATTAGCGGTGATCAGTGTTGTTG<br>TACCAGCTGTACACCGGAAGATTGTAGCGATCTGTGTAAAAATGGTAAACAGGCAGCACATGA<br>AGCCCAGAAACAGAAAAAATGTGCCAAATGTTGTAATGCCGGTGGTGAAAGCCATGAACTGT<br>GTTGTAGCATTGCAAGCGCAGGTATTGATTGTAATCCGTGTACCGCAGGTCTGCGTATGTGTTC<br>TTGGAGACCGT |
| SnTox3 | TAGGTCTCCAATGCTGGAACCTCGTGGTCCGGGTGACATCCAGCTGACCCGCGAAGAACACGA<br>AGCGATCTTCAACGGTTCCCCGTCTGACTGGACCGAAGACCCGAACTTCAAGCCGGACGTTCC<br>AGAACAGCAGCGCCCAGCGACCGCAAACGACCTGTCTAAACGTTACATCAAAGCCAACGACAT<br>CAACTTCGGTACGCGTTCTGTTCACGACTGCCGTGAGCGTACTGGTATTCAGCGTGACGTTAAA<br>GTTCGCGCGGACATCCCGTTCGAAACGGACGACGGTCCGAATCAGGTTCTGCGTGTAACCTGG<br>TCTAACGCCCTGAACGTTGACCGTTTTGACCCACTGCCGATCGTTACCGTTCCGGGTAACGCGG<br>CGTCTACCACCATCACCGCGATCCACGACTTCTGCCTGATGAACCCGACCACTTCTCCGCCGAC<br>CCGTTGCCTGTACCAGCTGCGTCAGCCGTTCACCCTGGGTTTCGACCGTACCCGTATGCACAAC<br>AACATCTACCTGACCCCTCCAAATCCGCAGCGTCCAACCATGCACGAAGTTTGCATCCGTGCGG<br>ACGAATGTCCAGCCGGTCGTGTTTTCCTGGAATGCTCTACTCGTACCTACGGTGCGATCCCGCG<br>TGGTGAATCTTGGAGACCGT |
| NIP2.1 | TAGGTCTCCAATGTACTATGTTGTCGTCTGCGTCCCCCGTGACGGCGCGGAAATCGGCGACGT<br>AGAGTGGGCCATCCAAAATCGCCGTCACGACCTGGCTCTGGGAGGTAAGGGGTTCTGGCGTG<br>GTCACAGCACTTCGTGCCATCGCAACGCAAACGCTGTGGTTGATGTTGTAGCTCTGTGTCGCTC<br>CGACCCATATATCGGAGCGCACCCAACGGTCTTAAAAGATGGGGCTTCTGTTTTGTGTCAGGC<br>ATCGGGAGCCCCAGACTGGCCGACTTGTACAGTGAATTGCTCTTGGAGACCGT |

**Supplementary Table S2 Continued. Genes used in this study**

| Name | Sequence |
|---|---|
| Human PDI | TACGTCTCACATATGGACGCACCTGAAGAGGAAGATCATGTTTTAGTCTTGCGTAAGTCTAACT<br>TCGCAGAGGCCTTGGCTGCTCATAAATACCTTTTGGTAGAGTTTTACGCACCTTGGTGCGGGCA<br>TTGCAAGGCTCTTGCTCCCGAATATGCGAAAGCCGCTGGAAAACTGAAGGCGGAAGGATCGG<br>AAATCCGCCTTGCCAAGGTGGATGCGACGGAGGAGTCAGACCTGGCCCAGCAATACGGGGTC<br>CGTGGGTACCCGACAATTAAATTCTTCCGCAATGGTGACACAGCATCACCGAAAGAATACACG<br>GCGGGGCGTGAGGCCGATGACATCGTTAACTGGCTTAAGAAACGCACTGGGCCAGCAGCGAC<br>TACCCTGCCAGATGGGGCAGCAGCTGAAAGTTTGGTGGAATCTTCCGAAGTGGCCGTAATTGG<br>GTTTTTTAAGGACGTAGAGTCAGACTCGGCAAAGCAATTCCTGCAAGCTGCTGAGGCAATTGA<br>CGATATTCCATTCGGTATCACCTCAAATTCGGACGTATTCTCTAAGTATCAGCTGGACAAAGAT<br>GGCGTAGTGTTATTTAAGAAATTTGATGAGGGTCGTAATAACTTCGAGGGCGAAGTGACCAAA<br>GAGAACCTTTTGGATTTTATTAAGCACAATCAACTGCCACTGGTTATCGAATTTACAGAACAGA<br>CGGCCCCTAAAATCTTCGGCGGTGAAATCAAGACCCACATTTTGCTGTTCCTGCCTAAGAGTGT<br>CTCGGACTACGATGGCAAACTGAGCAACTTTAAGACCGCGGCTGAGAGTTTTAAGGGGAAAAT<br>CCTTTTCATTTTCATCGACAGTGATCACACTGATAATCAACGCATTTTGGAGTTTTTCGGTCTGA<br>AAAAGGAAGAGTGCCCAGCAGTGCGTTTGATTACGCTTGAAGAAGAAATGACGAAATACAAG<br>CCTGAGTCTGAAGAATTGACCGCTGAGCGCATTACAGAGTTCTGTCATCGCTTCCTGGAGGGC<br>AAAATCAAACCTCACCTTATGTCGCAGGAATTACCAGAAGATTGGGATAAACAACCCGTCAAA<br>GTATTAGTGGGTAAAAATTTCGAAGATGTAGCGTTCGACGAAAAGAAGAACGTTTTCGTAGAG<br>TTCTATGCACCGTGGTGTGGTCACTGTAAGCAGCTGGCACCTATTTGGGATAAGCTGGGAGAA<br>ACGTATAAAGATCACGAAAATATCGTTATCGCGAAGATGGATTCAACTGCGAACGAAGTGGAA<br>GCGGTGAAGGTTCACTCTTTTCCAACACTTAAGTTTTTCCCTGCTTCGGCGGACCGTACAGTGA<br>TCGACTATAACGGCGAACGTACTCTTGATGGGTTTAAAAAATTTTTAGAGAGTGGTGGCCAGG<br>ATGGGGCAGGTGACGACGACGATCTTGAAGACTTAGAAGAAGCGGAGGAACCCGACATGGA<br>AGAAGATGACGACCAGAAAGCAGTAAAGGATGAATTGTGATAATCGAAGAGACGTA |
| Fol PDI | TACGTCTCACATATGGCAGATAGTGACGTCCACCAACTGACTAAGGACACGTTTGAGGAGTTT<br>GTTAAAAGTAACGACTTGGTACTTGCTGAATTTTTTGCTCCGTGGTGTGGTCATTGTAAAGCGT<br>TGGCACCCGAATATGAGGAAGCCGCTACCACCCTGAAGGAGAAGAATATCAAGCTGGCAAAG<br>ATCGACTGTACCGAAGAATCCGACCTGTGTAAGGATCAGGGGGTCGAGGGTTATCCCACTTTA<br>AAAGTGTTTCGCGGCTTGGAGAACGTCACGCCATACTCGGGGCAACGCAAGGCGGCGGGTAT<br>CACATCATATATGATTAAGCAGTCCTTACCTGCGGTATCAATTTTGACGAAGGATACACTTGAG<br>GAATTTAAAACTGCGGACAAGGTTGTTGTTGTAGCGTACTTAAATGCAGACGACAAGTCTAGC<br>AACGAAACTTTCTCTAAGTTAGCTGAAGGGTTACGTGACACGTATCTGTTCGGTGGGGTCAAT<br>GATGCTGCCGTCGCAGAAGCTGAGGGCGTTAAGGCGCCAGCTCTGGTTGTCTATAAATCGTTT<br>GATGAAGGTAAGAACACATTCACTGAGAAGTTCGAAGAAGATGCGATTGCCAGCTTTATCACC<br>ACCAGTGCAACGCCCTTGATCGGGGAAGTGGGTCCTGAAACATACGCTGGGTACATGTCGGC<br>GGGCATCCCCCTGGCCTACATCTTCTCAGAAACACCAGAAGAGCGTAAGGAACTGGGTGACGC<br>ATTGAAACCTATCGCGGAAAAGTTTAAAGGTAAAATTAATTTCGCAACTATTGACGCGAAGGC<br>TTTCGGGGCACACGCGGGGAATTTAAACTTGAAGGCCGACAAGTTTCCTTCGTTTGCGATTCAA<br>GAAGTAGTCAAAAACCAGAAATTTCCGTTTGATCAAGAGAAGGAGATCACCCATGACAACATT<br>GCTAAGTTCGTGGAAGATTTTGCGGCTGGAAAGATCGAACCCTCCATTAAGTCTGAGCCTATTC<br>CTGAAACGCAGGAGGGACCAGTCACGGTAGTCGTGGCCAAATCGTACAACGACATCGTACTTG<br>ATGACACGAAAGACGTGTTGATCGAATTCTACGCTCCCTGGTGCGGCCACTGTAAGGCATTGG<br>CACCCAAATACGAGGACTTGGCGTCACAATTTGCCGCGTCCGAGTTCAAAGATAAGGTAGTCA<br>TCGCCAAGGTGGACGCAACCCTTAACGATGTACCTGATGAGATCCAGGGATTCCCTACAATCA<br>AGTTGTATGCAGCAGGAGCCAAAGACGCACCGGTCACATACCAAGGTTCACGCACCGTCGAA<br>GATTTGGCCAATTTCATTAAAGAGAACGGCAAATACAAGGCTGAGTTGCCTGTAAAGGAGGAG<br>GGGACTGAGGAGGCCGCGCCCGCTGCGAGTGAAGAAAAGAAGGAGGAAAAGAAGGATGCG<br>GAAGAAGAAGACGTCCATGACGAATTGTGATAATCGAAGAGACGTA |

**Supplementary Table S2 Continued. Genes used in this study**

| Name | Sequence |
|---|---|
| Erv1p | TACGTCTCACATATGAAGGCTATTGACAAGATGACGGACAACCCGCCGCAAGAAGGGTTATCAGGTCGTAAAATCATTTACGACGAGGACGGCAAGCCTTGTCGTTCTTGTAATACTCTTTTGGACTTTCAGTACGTTACCGGCAAAATCAGTAACGGCTTAAAGAATCTTAGCTCGAACGGGAAGTTGGCTGGGACCGGGGCTCTTACGGGTGAAGCATCCGAGTTAATGCCAGGGAGCCGCACGTACCGTAAAGTAGACCCGCCGGACGTAGAGCAGTTAGGCCGTTCATCATGGACTTTGTTACATTCTGTCGCCGCATCGTACCCTGCACAGCCTACCGATCAGCAAAAGGGAGAGATGAAGCAATTTTTAAATATCTTTTCTCACATTTACCCGTGCAATTGGTGCGCGAAAGACTTTGAAAAATACATTCGTGAAAACGCGCCACAAGTCGAGTCACGTGAGGAGTTAGGTCGTTGGATGTGCGAGGCCCACAACAAAGTCAACAAAAAGTTACGTAAGCCGAAGTTCGACTGTAATTTCTGGGAAAAGCGCTGGAAGGATGGCTGGGACGAGTGATAATCGAAGAGACGTA |
| FolErv2 | TACGTCTCACATATGAGCGGCCCGAGTCGTTCAATCCTTCCCAAATACCAGAACGATGTAGAGTTACCGCTTAAGGAAGCCCCACGCTCGGAGTTCGCCGCTGACCTTGGTGCTCTTCCGGCCGGTTTGTTGGATGGGAACTCGATTGCTCCAAAATTGGAGAACGCAACACTTAAGGCGGAGTTGGGGCATGCGACTTGGAAATTCTTACACACGATGATGGCCCGCTTCCCTGACAAACCGACCAAAGATGACCGCATGGCACTTGAAACCTTTATGCACTTATTCGCCCGCTTATATCCTTGCGGTCAATGCGCGGCTCACTTTCAAAAGTTACTGGCACAATACCCTCCACAAACATCATCCCGCAATGCCGCGGCCGGATGGTTGTGCTTTGCTCATAATATCGTCAATGAACGCGTCCACAAACCGTTATTCGATTGCGAGAATATCGGGGACTTCTACGATTGCGGGTGCGGCGACAAAGACAAGAAGGAGGGTGGAGCAGAAACCACTGCAGCCGCGCAAGCCGAAGGCCCAGACCAGGAGCTTCATAAACACTAATCGAAGAGACGTA |
| FOXG_09255* | MARRQHLTLTFILVLGVFFTLSYFFSGPSRSILPKYQNDVELPLKEAPRSEFAADLGALPAGLLDGNSIAPKLENATLKAELGHATWKFLHTMMARFPDKPTKDDRMALETFMHLFARLYPCGQCAAHFQKLLAQYPPQTSSRNAAAGWLCFAHNIVNERVHKPLFDCENIGDFYDCGCGDKDKKEGGAETTAAAQAEGPDQELHKH |

*Amino acid sequence of the correctly annotated FOXG_09255 gene based on RNAseq data.
